# Photoacoustic imaging for the prediction and assessment of response to radiotherapy *in vivo*

**DOI:** 10.1101/329516

**Authors:** Márcia Martinho Costa, Anant Shah, Ian Rivens, Carol Box, Tuathan O’Shea, Jeffrey Bamber, Gail ter Haar

## Abstract

Radiotherapy is commonly used for cancer therapy, although its efficacy is reduced in hypoxic regions of tumours. Photoacoustic imaging (PAI) is an emergent, non-invasive imaging technique that allows the measurement of blood oxygen saturation (sO_2_) which inversely correlates with hypoxia in tissue. The potential use of PAI as a prognostic tool for radiotherapy outcome was investigated in a head and neck cancer model *in vivo*. PAI was performed before delivering a single fraction (10, 20 or 30 Gy) treatment. The results show that tumours with pre-treatment higher blood sO_2_ responded better than those with lower levels in the 10 and 20 Gy groups. For the 30 Gy group, treatment response was independent of blood sO_2_. The haemoglobin content of the tumours was not correlated with their response to any of the radiation doses studied. Changes in sO_2_, monitored at 24 h and 96 h following 10 and 20 Gy doses, showed that tumours that were subsequently unresponsive to treatment had an increase in blood sO_2_ at both time points compared to those which subsequently regressed after radiotherapy. The results suggest that sO_2_ values measured by photoacoustic imaging can be used before, and shortly after, irradiation to predict subsequent treatment response.

## Introduction

Radiotherapy (RT) is one of the most effective cancer treatments. However, tumour cells are often hypoxic, making them resistant to radiation damage [1-4] and, consequently, the probability of disease recurrence after treatment increases [5, 6]. The use of higher radiation doses can reduce this problem, but this increases morbidity due to damage to normal tissue and late radiation side effects [7]. The distribution of hypoxic regions within a tumour is known to be both spatially heterogeneous and temporally variant [8, 9], over periods from hours to days. Identification of hypoxic tumours or hypoxic regions of a tumour may identify patients or tumour regions that would require dose escalation, potentially sparing the patients from ineffective therapy.

The distribution of hypoxia can be inferred using a number of imaging approaches [10], some of which involve the use of hypoxia reporter molecules or quantification of blood oxygen saturation (sO_2_). Positron Emission Tomography (PET) is the current gold standard, but the poor perfusion restricts access of the hypoxia-targeted imaging probes to map the spatial heterogeneity in hypoxia [11, 12]. This is also true for phosphorescence quenching [13], electron paramagnetic resonance [14], dynamic contrast enhanced-magnetic resonance imaging (DCE-MRI), and in general for such tracer based imaging approaches.

Tumour hypoxia has been shown to be associated with a decrease in blood sO_2_ [15]. Blood-oxygen level dependent MRI (BOLD-MRI) can infer the hypoxia distribution in tumour tissue by imaging the transition of deoxyhaemoglobin (Hb) to oxyhaemoglobin (HbO_2_) in blood, on changing from air (21% oxygen) to 100% oxygen gas breathing [16-20]. However, BOLD-MRI cannot estimate absolute blood sO_2_.

Photoacoustic imaging (PAI) has demonstrated the potential for quantifying absolute blood sO_2_ by identifying the unique optical absorption signatures of Hb and HbO_2_ without the need for tracers [21]. Light, delivered by a laser, is absorbed by components of tissue, which undergo thermoelastic expansion, thus stimulating transient wideband ultrasonic wave generation. The intensity (or amplitude) of acoustic signals is proportional to the absorption coefficient and the optical fluence [21] of the tissue, and can be measured on the tissue surface using ultrasound detectors.

PAI has been used to identify regions of tissue more likely to be hypoxic, pre-clinically in subcutaneous tumours and in the brain [19, 22-26], and clinically in breast cancer patients [27]. This was achieved by comparing the results obtained from photoacoustic imaging with immunohistochemical hypoxia markers and other imaging modalities, such as BOLD-MRI and microbubble contrast-enhanced high frequency ultrasound. These studies found good correlation between regions with low sO_2_ and those identified as potentially hypoxic. Our group has also found a good correlation between the distribution of pimonidazole, an exogenous hypoxia marker, and the sO_2_ distribution measured by PAI [28].

Here, we hypothesize that PAI can be used for assessing tumour oxygenation status in a preclinical head and neck cancer model before, and after, single fraction radiotherapy treatments, and that these blood sO_2_ measurements would correlate with treatment outcome. We have also also explored the capability of individual haemoglobin components (Hb, HbO_2_ and total haemoglobin, HbT) and ΔsO_2_ (sO_2_ (oxygen) - sO_2_ (air)) to predict and monitor radiotherapy response. The last parameter can be obtained from an oxygen challenge test and it has been shown to allow the delineation between different tumours’ vascular characteristics [29], as oxygen can be used as a contrast agent for hypoxia studies.

## Results

### Pre-irradiation imaging for predicting tumour outcome

We assessed the potential of PAI to predict RT efficacy by imaging animals bearing subcutaneous CAL^R^ tumours, 24 h and immediately before delivering a single boost RT dose (10, 20 or 30 Gy) to the tumour. Data was acquired under both medical air and 100%-oxygen breathing conditions. One imaging dataset acquired prior to delivering a RT dose of 10 Gy was excluded due to an artefact masking the signal in the tumour (Fig. S1).

Fig. 1 shows the normalised tumour growth curves (caliper measurements) for the control (CTRL), 10, 20 and 30 Gy cohorts, subdivided into groups by the level of response to RT dose: full-responders (FR), partial-responders (PR) and no-responders (NR). The NR growth curves, shown in red, are within the range of variability of the tumour growth for the control cohort, shown in black. There was no difference in the survival of the NR cohort in comparison to the control group. FR, represented in green, had a tumour volume decrease, reaching 0 mm^3^, i.e. no macroscopic signs of disease were palpable. The lack of palpable tumour mass was maintained for the remaining 60 day follow-up period. The average time after-RT at which tumour volume, V, decreased below the initial volume, V_0_, was 29±8 days for 10 Gy, 9±6 days for 20 Gy and 11±5 days for 30 Gy. The survival of the full responders was increased compared to the controls. The PR growth curves, represented in blue, showed that there was an increase in the survival of the mice compared to that of the control cohort, i.e. some tumours regrew and some maintained a stable volume above V_0_.

**Figure 1:**
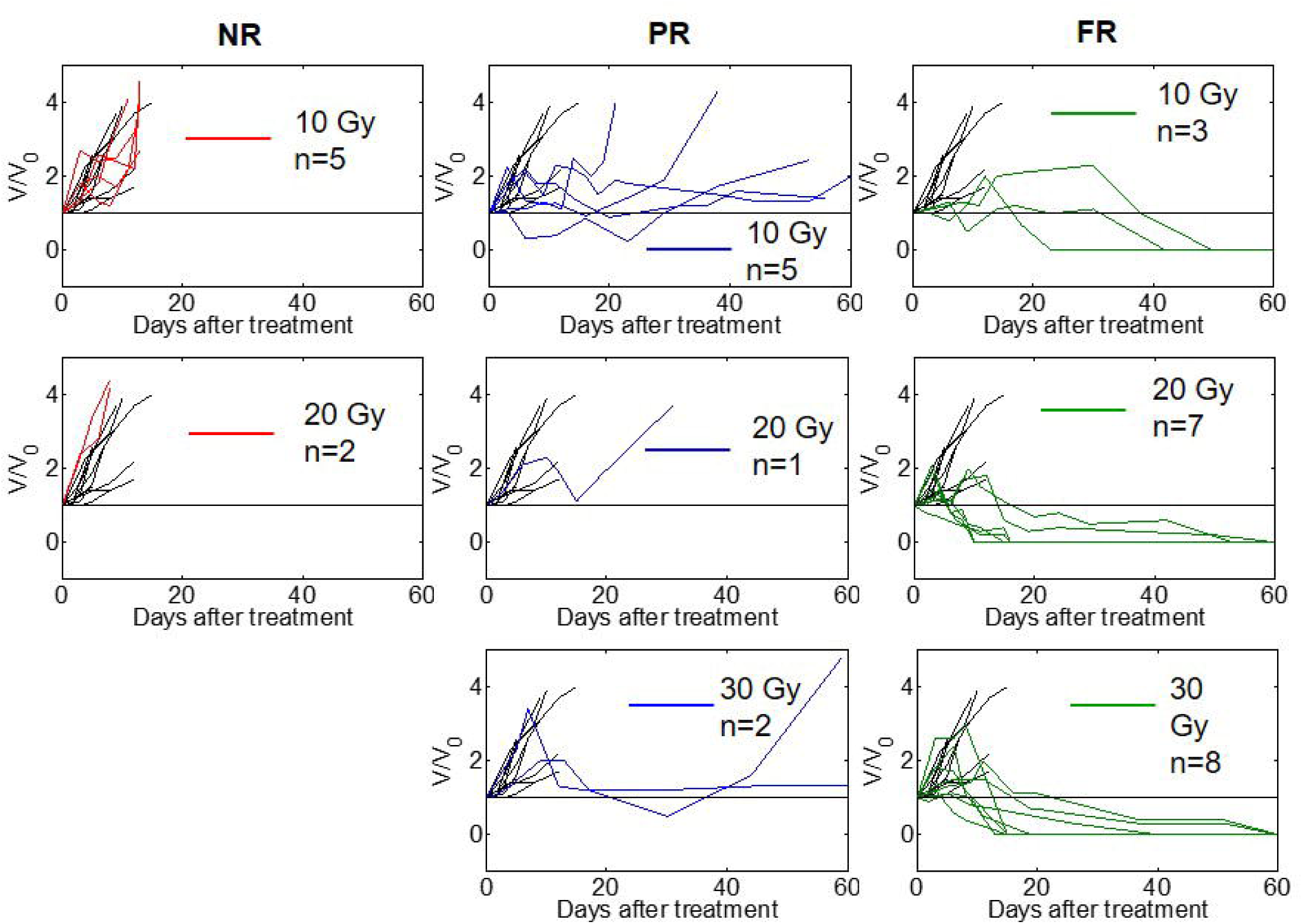
Normalised tumour growth curves for controls (black lines), 10, 20 and 30 Gy irradiated cohorts, subdivided as no-responders (NR), partial-responders (PR) or full-responders (FR). Tumour volume measured at each time point after treatment (V) was normalised to V_0_ the volume on treatment day. Mean tumour volumes at onset of treatment (V_0_) were: 10 Gy, 201±15 mm^3^; 20 Gy, 216±30 mm^3^; 30 Gy, 214±19 mm^3^.

#### Average blood sO_2_ results

Fig. 2A shows representative ‘oxymaps’, i.e. maps of the calculated blood sO_2_, for the tumour central slice, for the 10 Gy cohort, from the FR, PR and NR groups, immediately pre-RT, during air-breathing. The ‘oxymaps’ are shown overlaid on greyscale photoacoustic images, in order to visualise where, within the tumour, the blood sO_2_ is being calculated. ‘Oxymaps’ of tumours of FRs had majority of pixels corresponding to high values of sO_2_, while those of PR and NR tumours had substantial pixels corresponding to low blood sO_2_. The ‘oxymaps’ for the entire 10 Gy cohort, immediately pre-RT (air-breathing), are shown in Fig. S1.

**Figure 2:**
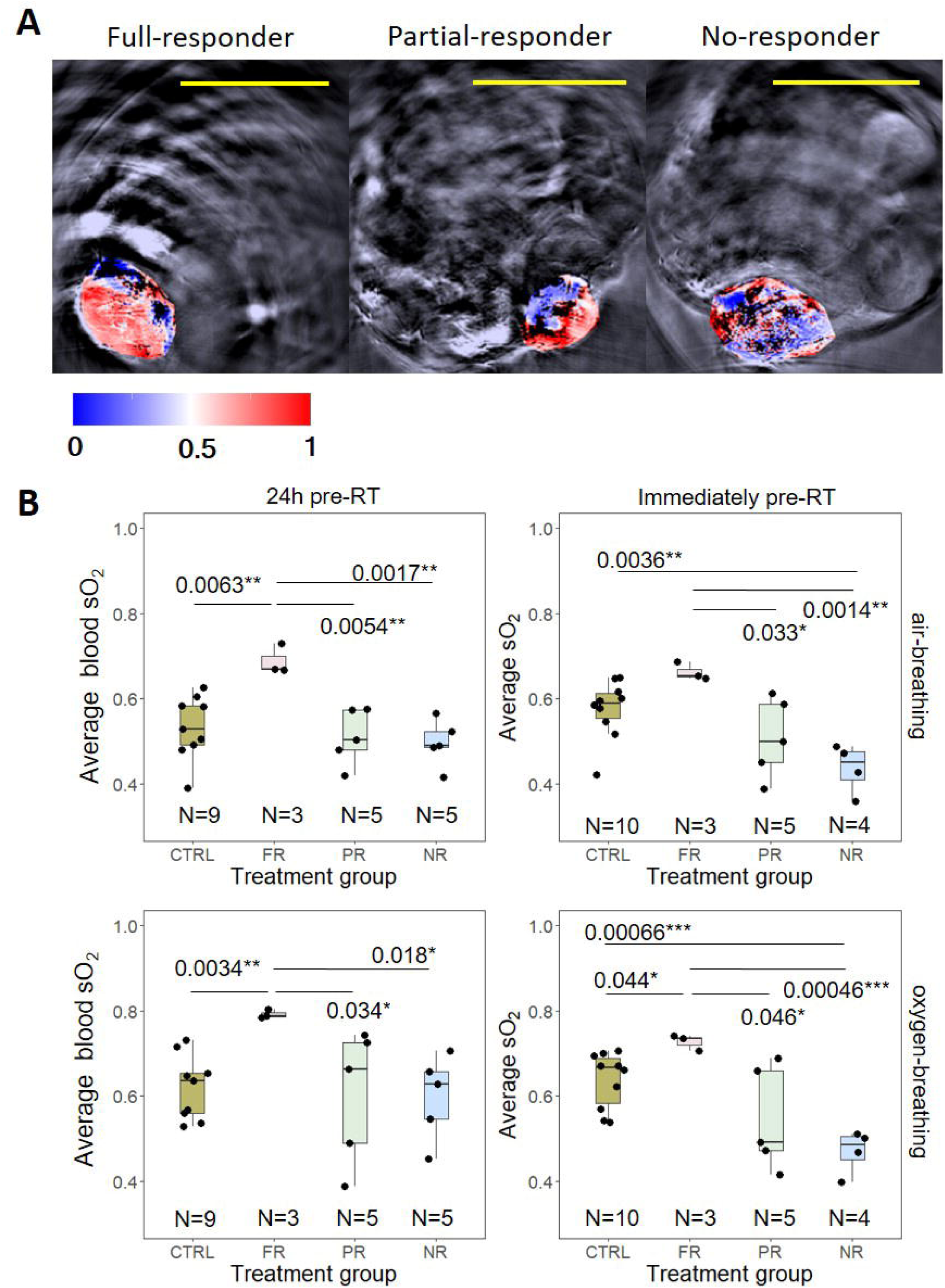
**A:** Three examples of ‘oxymaps’ obtained pre-RT for the 10 Gy cohort, during air-breathing, for FR, and NR tumours. The ‘oxymaps’ show the calculated blood sO_2_ pixel-by-pixel and are overlaid in a greyscale photoacoustic image. Blue regions represent blood sO_2_ below 0.5 and pink-to-red regions blood sO_2_ above 0.5. Black regions represent pixels in which the calculated total amount of haemoglobin falls below 0, i.e. the amount of haemoglobin was below the system’s noise threshold. Yellow bars = 10 mm; B: body of the mouse; W: water, used as an acoustic coupling medium. **B:** Average blood sO_2_ measured 24 h and immediately before delivering 10 Gy irradiation. Black circles represent the average of three central PAI slices for each tumour. Black horizontal lines in the boxplots represent the median sO_2_ with the box extremities representing the 25% and 75% percentiles and the whiskers representing the maximum and minimum values. Full responders (FR, n=3) had a statistically significantly higher average blood sO_2_ than partial-(PR, n=5) and no-responders (NR, n=5) during both air-and oxygen-breathing, both 24 h and immediately pre-RT. * p-value < 0.05; ** p-value < 0.01; *** p-value < 0.001 using a paired Student’s t–test.

The blood sO_2_ values were averaged over three central tumour slices and measured 24 h and immediately before delivering 10 Gy irradiation. As seen in Fig. 2B and Table 1, the FR cohort had statistically significantly higher levels of blood sO_2_ (0.66±0.02 and 0.73±0.02 for air-and oxygen-breathing, respectively, immediately pre-RT) in comparison to the PR (0.51±0.08 and 0.55±0.1 for air-and oxygen-breathing, respectively) and NR (0.44±0.05 and 0.47±0.04 for air-and oxygen-breathing, respectively) cohorts. FRs had a significantly higher average blood sO_2_ values compared to the control group, apart from immediately pre-RT measurements during air-breathing (Table 1). On the other hand, NR tumours, imaged immediately pre-RT, had significantly lower average blood sO_2_ (0.44±0.05 and 0.47±0.04 for air-and oxygen-breathing, respectively) than the controls (0.58±0.06 and 0.64±0.06 for air-and oxygen-breathing, respectively).

**Table 1:** Average blood sO_2_ (± standard deviation, s.d.), acquired by PAI, for tumours irradiated with 0, 10, 20 or 30 Gy. Treatment response groups are: FR (full-responders); PR (partial-responders) and NR (no-responders) and images were acquired during both air-and oxygen-breathing 24 h pre-and immediately pre-RT. Statistically significant differences between FR and the other response groups are indicated by: * p-value < 0.05; ** p-value < 0.01; *** p-value < 0.001 using a paired Student’s t–test.

| | | Average blood sO<sub>2</sub> (± s.d.) for each treatment response | | | |
| --- | --- | --- | --- | --- | --- |
|  |  | Air-breathing |  | Oxygen-breathing |  |
| Dose (Gy) | Treatment response | 24 h pre-RT | Immediately pre-RT | 24 h pre-RT | Immediately pre-RT |
| 0 | Control (n=10) | 0.52 ± 0.07 | 0.58 ± 0.06 | 0.62 ± 0.07 | 0.64 ± 0.06 |
| 10 | FR (n=3) | 0.69 ± 0.03 | 0.66 ± 0.02 | 0.79 ± 0.01 | 0.73 ± 0.02 |
| | PR (n=5) | 0.51 ± 0.06** | 0.51 ± 0.08* | 0.60 ± 0.14* | 0.55 ± 0.1* |
| | NR (n=5) | 0.50 ± 0.05** | 0.44 ± 0.05** | 0.60 ± 0.09* | 0.47 ± 0.04*** |
| 20 | FR (n=7) | 0.63 ± 0.08 | 0.63 ± 0.09 | 0.73 ± 0.07 | 0.71 ± 0.11 |
| | PR+NR (n=3) | 0.45 ± 0.02** | 0.50 ± 0.02* | 0.57 ± 0.03* | 0.55 ± 0.01* |
| 30 | FR (N=8) | 0.55 ± 0.06 | 0.56 ± 0.07 | 0.63 ± 0.12 | 0.61 ± 0.10 |
| | PR (n=2) | 0.48 ± 0.06 | 0.45 ± 0.07 | 0.48 ± 0.06 | 0.50 ± 0.03 |

The 20 Gy group had only one PR with average blood sO_2_ levels similar to those of NR. Therefore all the PAI data for this animal was combined with the NR data, and this combined group was compared with the FR cohort. The tumour averaged blood sO_2_ values, obtained 24 h and immediately before delivering 20 Gy irradiation, are shown in Fig. 3 and Table 1. During both air-and oxygen-breathing imaging conditions, tumours which went on to have a full response (n=7) had statistically significantly higher blood sO_2_ (0.63±0.09 and 0.71±0.11 for air-and oxygen-breathing, respectively, immediately pre-RT) than those with partial-and no-response (n=3, 0.50±0.02 and 0.55±0.01 for air-and oxygen-breathing, respectively, immediately pre-RT). This difference was also observed 24 hours pre-RT. Twenty-four hours pre-RT, FR had significantly higher average blood sO_2_ (0.63±0.08 and 0.73±0.07 for air-and oxygen-breathing) than the control cohort (0.52±0.07 and 0.62±0.07 for air-and oxygen-breathing). NR tumours had significantly lower sO_2_ immediately pre-RT compared to the control cohort, but only during oxygen-breathing (0.55±0.01 and 0.64±0.06, for NR and control cohorts, respectively).

**Figure 3:**
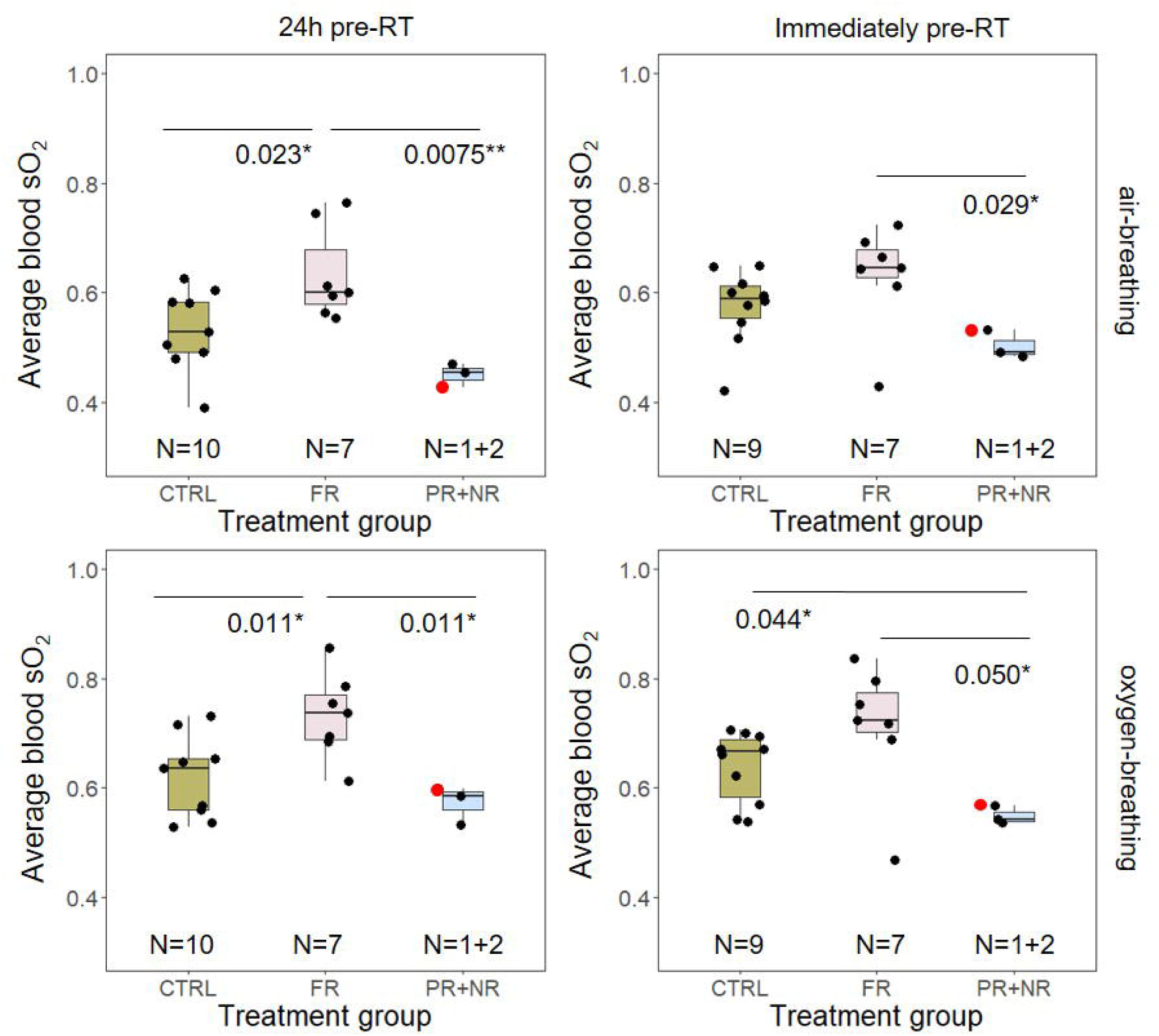
Average blood sO_2_ measured 24 h and immediately before delivering 20 Gy irradiation. Black circles represent the average of three central PAI slices for each tumour. Red circle represents the partial-responder. Black horizontal lines in the boxplots represent the median average, with the box extremities representing the 25% and 75% percentiles and the whiskers representing the maximum and minimum values. Full responders (FR, n=7) had a statistically significantly higher average blood sO_2_ than combined partial-(PR, n=1) and no-responders (NR, n=2) during both air-and oxygen-breathing, both 24 h and immediately pre-RT. * p-value < 0.05; ** p-value < 0.01; *** p-value < 0.001 using a paired Student’s t –test.

For tumours irradiated with 30 Gy the association between pre-treatment average blood sO_2_ and radiation response was weaker. This could be because majority of the tumours responded to the high dose, irrespective of their blood sO_2_. One tumour had a low average blood sO_2_ (0.35±0.01 24 h pre-RT and 0.41±0.01 immediately pre-RT) and went on to respond fully to the radiation (Fig. 4 and Table 1). Although the average blood sO_2_ for FR was higher (0.56±0.07 and 0.61±0.10 for air-and oxygen-breathing, immediately pre-RT) than for PR (0.45±0.07 and 0.50±0.03 for air-and oxygen-breathing, immediately pre-RT), there were no statistically significant differences between the two groups. Nevertheless, PR had significantly lower average blood sO_2_ than the control cohort (Table 1).

**Figure 4:**
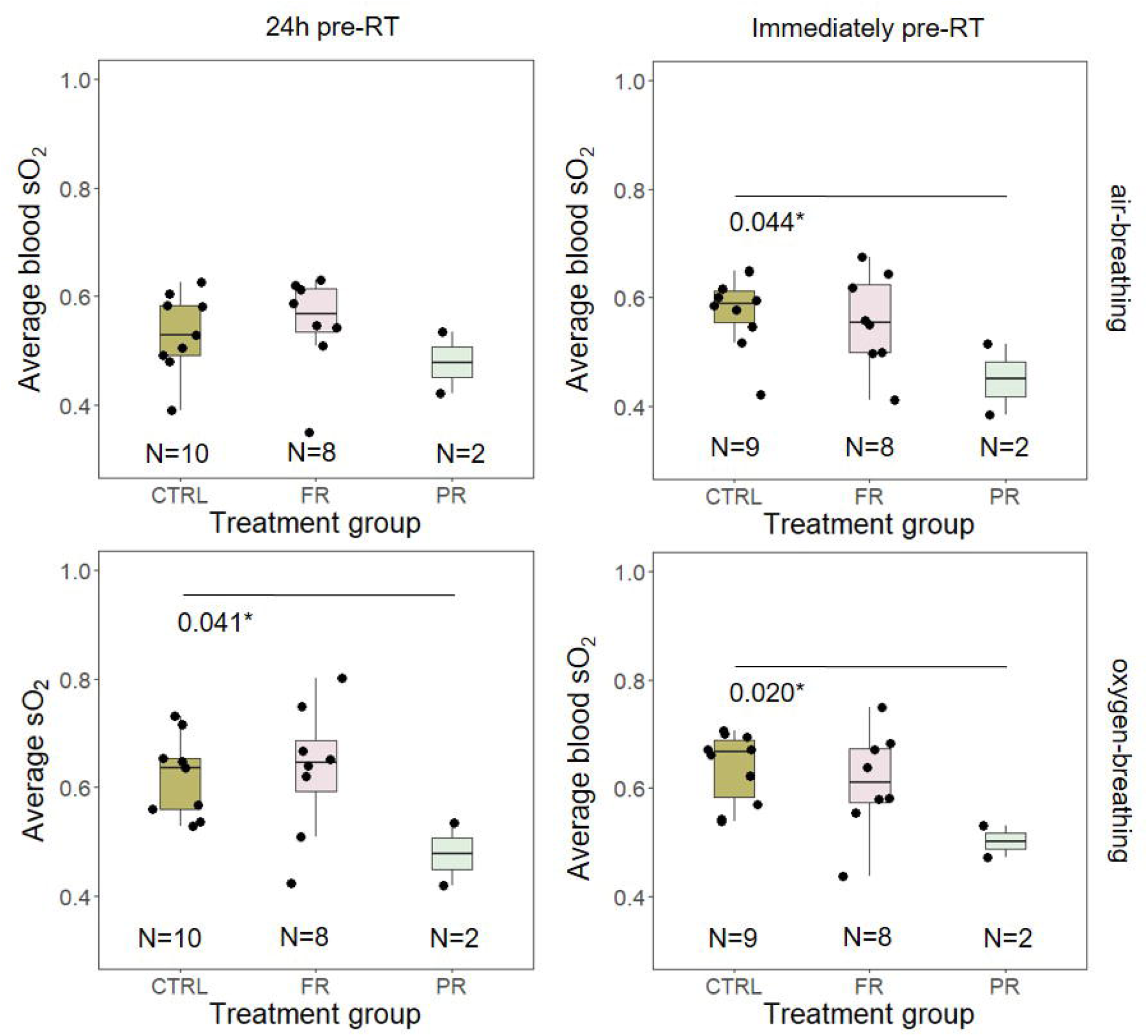
Average blood sO_2_ measured 24 h and immediately before delivering 30 Gy irradiation. Black circles represent the average of three central PAI slices for each tumour. Black horizontal lines in the boxplots represent the median average, with the box extremities representing the 25% and 75% percentiles and the whiskers representing the maximum and minimum values per group. No statistical differences (Student’s t-test) were observed between full-(FR, n=8) and partial-responders (PR, n=2).

#### *ΔsO*_*2*_ *and haemoglobin results*

The difference in measured blood sO_2_ during air-and oxygen-breathing (ΔsO_2_) and its relationship with treatment outcome was also investigated, 24h and immediately pre-RT. No significant differences were found between FR, PR and NR groups, at 10, 20 or 30 Gy (Tables S1-S3).

The average oxy-(HbO_2_), deoxy-(Hb) and total haemoglobin (HbT) were measured for each tumour, at the two time points before treatment, to study if haemoglobin levels could also be used as a predictive factor to radiotherapy outcome. Mostly, there were no statistically significant differences in these PAI parameters between the FR group and PR/NR irrespective of the radiation dose (Tables S1-S3). However, for tumours irradiated with 10 Gy, there was a significant difference between FR and PR groups 24 h pre-RT (p-value of 0.041* during air-breathing and 0.0019** for oxygen-breathing; Table S1). FR had, on average, less HbO_2_ (63 ± 17 A.U. for air-breathing, 54 ± 17 A.U. for oxygen-breathing) than PR (143 ± 44 A.U. and 120 ± 17 A.U. for air-and oxygen-breathing, respectively). This significance was only observed for this comparison, so it is likely to be an outlier.

### Post-irradiation imaging for treatment monitoring

The potential of PAI to monitor RT treatment response was investigated by estimating blood sO_2_ and haemoglobin levels of animal tumours 24 h and 96 h post-RT.

#### *Average blood sO*_*2*_ *results*

Fig.5A shows representative tumour ‘oxymaps’, for the 10 Gy cohort, obtained 96 h post-RT (air-breathing), for one FR and one NR. Immediately pre-RT, the FR ‘oxymaps’ had predominantly red pixels, indicating an average sO_2_ > 0.5, while NR ‘oxymaps’ had largely white and blue pixels, suggesting sO_2_ ≤ 0.5. Ninety-six hours post-RT, an increase in blue pixels at the centre of the FR tumour (Fig. 5), was observed, suggesting a decrease in the mean blood sO_2_. On the other hand, in the NR tumour, an increase in the red pixels or high blood sO_2_ regions was observed 96 h post-RT, particularly in the margins of the tumour. The remaining ‘oxymaps’ for the 10 Gy cohort are shown in Fig. S1

**Figure 5:**
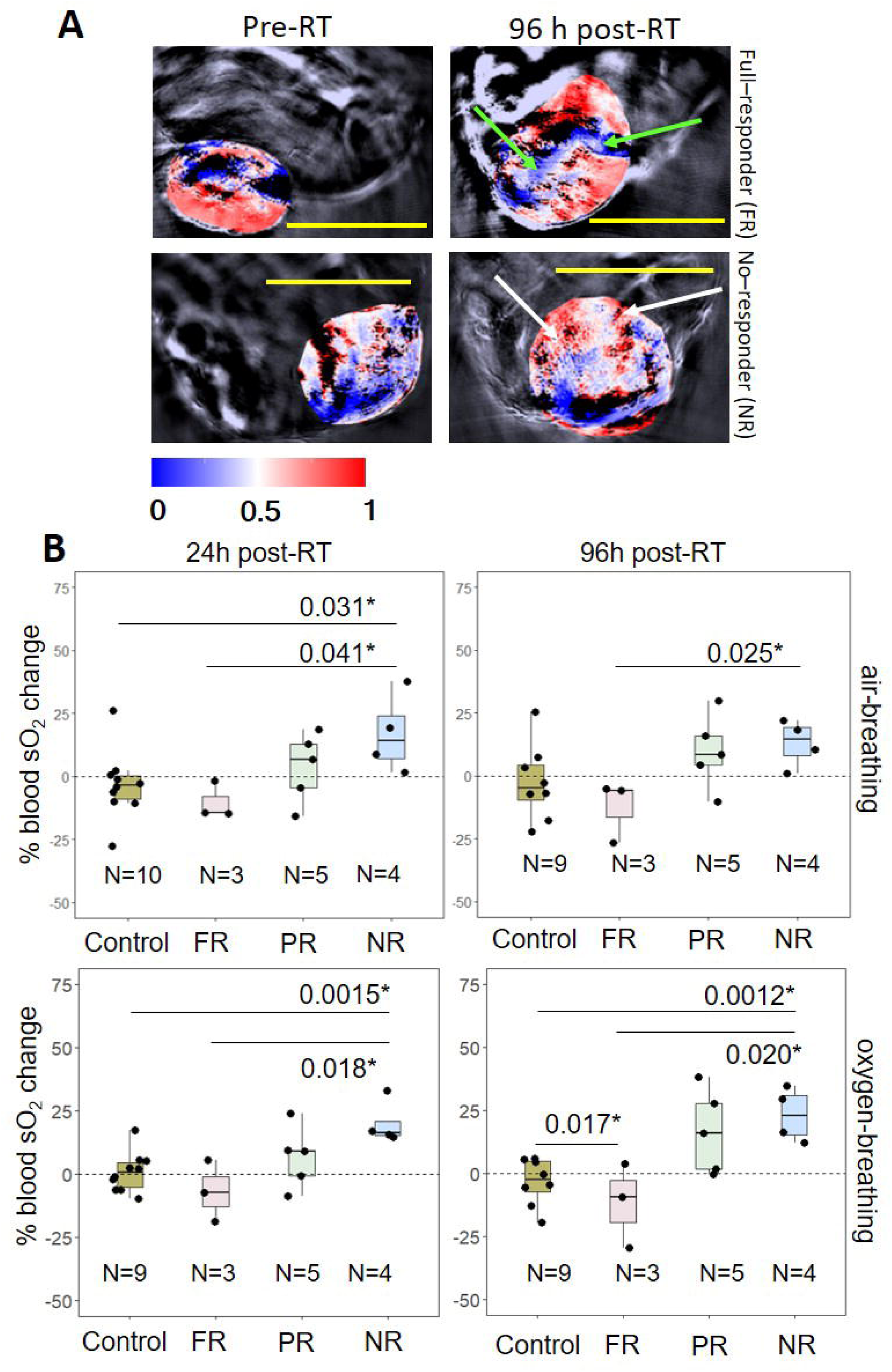
**A:** Examples of tumour ‘oxymaps’ obtained immediately pre-RT and 96 h post-RT for 10 Gy cohort, showing the differences in blood sO_2_ between FR and NR at the two time points (air-breathing). Green arrows indicate blue regions (low blood sO_2_) observed 96 h post-RT for the FR, which were not present pre-RT. White arrows indicate red regions, i.e. regions with high blood sO_2_, for the NR 96 h post-RT, which were not observed pre-RT. Yellow bar = 10 mm. **B:** Percentage change in average blood sO_2_ for the 10 Gy RT group between 24 hours, left column, and 96 hours post-RT, right column, and immediately pre-irradiation. Black circles represent the change in average blood sO_2_ for each tumour measured over 3 central slices. Black horizontal lines in the boxplots represent the median average, with the box extremities representing the 25% and 75% percentiles and the whiskers representing the maximum and minimum values per group. FR = full-responders; PR = partial-responders; NR = no-responders. * p-value < 0.05; ** p-value < 0.01 using a paired Student’s t-test.

The percentage change in average blood sO_2_ measured at 24 h and 96 h post-RT with respect to the baseline (i.e. that measured for the same tumour immediately pre-RT) for the control and 10 Gy cohorts are shown in Fig. 5B and Table 2. The mean percentage change in average blood sO_2_ for the control animals was close to zero, for both air-and oxygen-breathing, at both time points. As seen in Fig. 5 the mean percentage change in blood sO_2_ for the FR and the NR cohort, was consistently negative and positive, respectively at both time-points post-RT. There was no statistically significant difference between the control and FR cohorts. Interestingly, the NR group showed a statistically significant increase in blood sO_2,_ at both 24 h and 96 h post-RT, in comparison to either the FR or control groups, during air-or oxygen-breathing imaging (Table 2). The partial-responders (PR) had a positive percentage change in average sO_2_ at both 24 h and 96 h post-RT, although there were no statistically significant differences between these and the remaining response groups or control cohort.

**Table 2:** Summary of the % change from pre-treatment baseline of average blood sO_2_ measured 24 h and 96 h post-RT, per treatment response group, for control, 10, 20 and 30 Gy cohorts, for both air-and oxygen-breathing. Statistical differences between PR and NR to the other response groups are indicated by: * p-value < 0.05; ** p-value < 0.01.

| Dose<br>(Gy) | Treatment<br>response | % change in average blood sO<sub>2</sub> (± s.d.) |  |  |  |
| --- | --- | --- | --- | --- | --- |
|  |  | Air-breathing |  | Oxygen-breathing |  |
|  |  | 24 h post-RT | 96 h post-RT | 24 h post-RT | 96 h post-RT |
| 0 | Controls | -3 ± 13 | -3 ± 14 | 0.8 ± 7 | -3 ± 9 |
| 10 | FR (N=3) | -10 ± 6 | -13 ± 10 | -7 ± 10 | -12 ± 14 |
|  | PR (N=5) | 4 ± 12 | 10 ± 13 | 7 ± 11 | 17 ± 15 |
|  | NR (N=5) | 17 ± 14 | 13 ± 8* | 20 ± 7* | 23 ± 9** |
| 20 | FR (N=7) | 0.8 ± 9 | -8 ± 18 | 0.3 ± 5 | -6 ± 21 |
|  | PR+NR<br>(N=3) | -10 ± 5 | 25 ± 16* | -9 ± 4 | 26 ± 15* |
| 30 | FR (N=8) | 6 ± 17 | 7 ± 18 | 6 ± 11 | 13 ± 18 |
|  | PR (N=2) | 19 ± 5 | 21 ± 10 | 15 ± 2* | 20 ± 4* |

The percentage change in blood sO_2_ at 24 and 96 h post 20 Gy irradiation, with respect to blood sO_2_ pre-RT, is shown in Fig. 6 and Table 2. Similar to the observations in the 10 Gy cohort, the FRs and NRs had a mean negative and positive percentage change in blood sO_2_, respectively, for both time points post RT. There was no statistical difference between the FR and NR groups for the 24 hour time-point. However, at 96 h post RT, there was a statistically significant difference in the percentage change of blood sO_2_ for the NRs (25 ± 16 % for air-breathing and 26 ± 15% for oxygen-breathing) in comparison to the FRs (-8 ± 18 % for air-breathing and -6 ± 21% for oxygen-breathing). The ‘oxymaps’ of the 20 Gy cohort are shown in Fig. S2.

**Figure 6:**
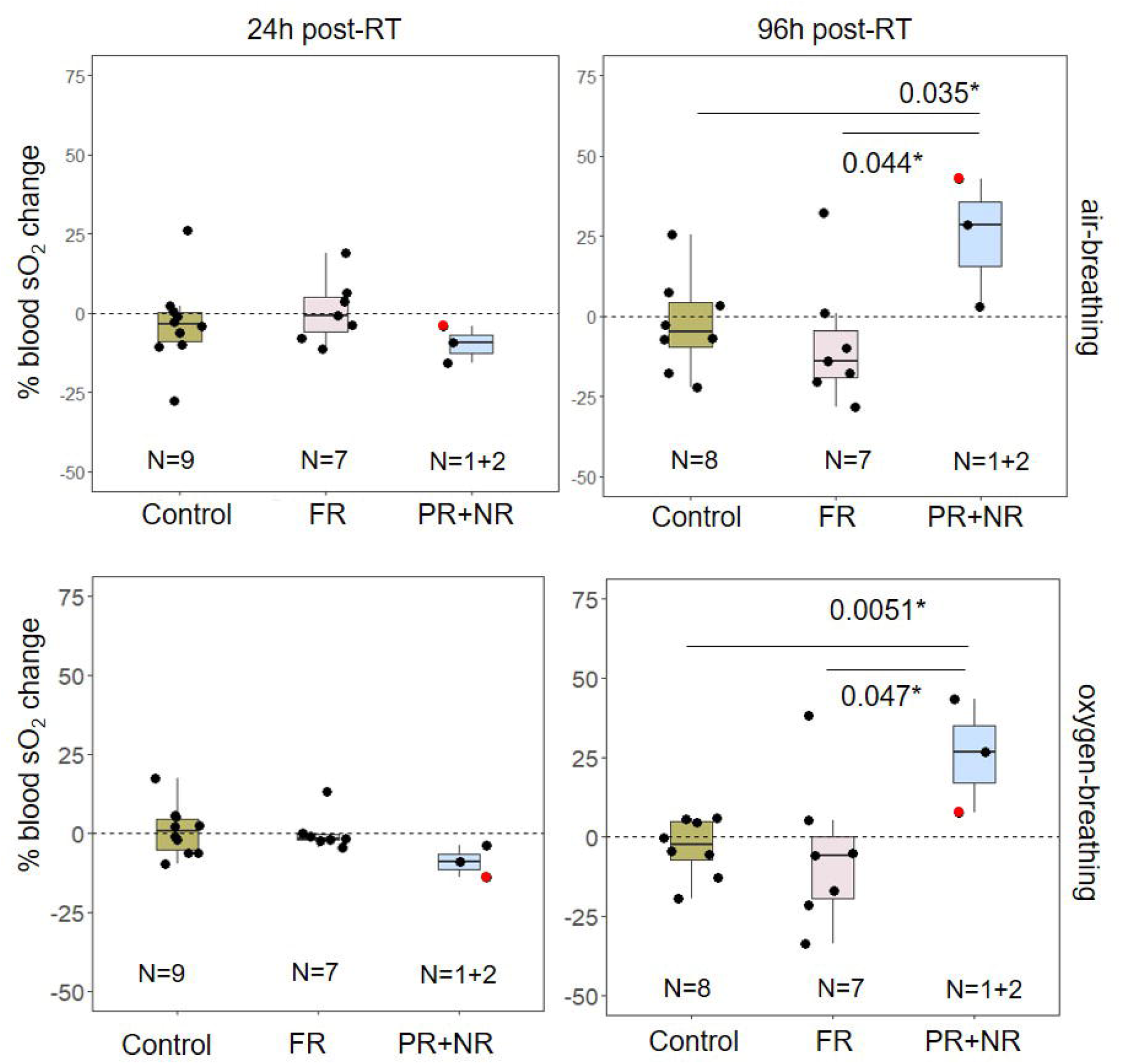
Percentage change in average blood sO_2_ for the 20 Gy RT group between 24 hours, left column, and 96 hours post-RT, right column, and immediately pre-irradiation. Black circles represent the change in average blood sO_2_ for each tumour measured over 3 central slices. Red circle corresponds to the partial-responder. Black horizontal lines within the boxplots represent median blood sO_2_ change per group, with the box extremities representing the 25% and 75% percentiles and the whiskers representing the maximum and minimum values per group. FR = full-responders; PR+NR = partial-and no-responders. * p-value < 0.05; ** p-value < 0.01 using a paired Student’s t-test.

‘Oxymaps’ of tumours (Fig. 7) depict changes in blood sO_2_ which occurred 96 h post-irradiation with 30 Gy, during air breathing. One of the full-responders had high blood sO_2_ levels throughout most of the tumour prior to irradiation (Fig. 7, top row), which was greatly reduced by 96 h post-RT. A different tumour, which also responded fully to treatment, had a large region with low baseline blood sO_2_, which displayed increased oxygen saturation levels 96 h post-RT, particularly in the middle and at the margins closest to the mouse body. The ‘oxymaps’ for the PR tumour also showed regions with increased sO_2_ (pink) 96 h post-RT compared with the baseline. The remaining ‘oxymaps’ for the 30 Gy cohort, pre-RT and 96 h post-RT (air-breathing) are shown in Fig. S3.

**Figure 7:**
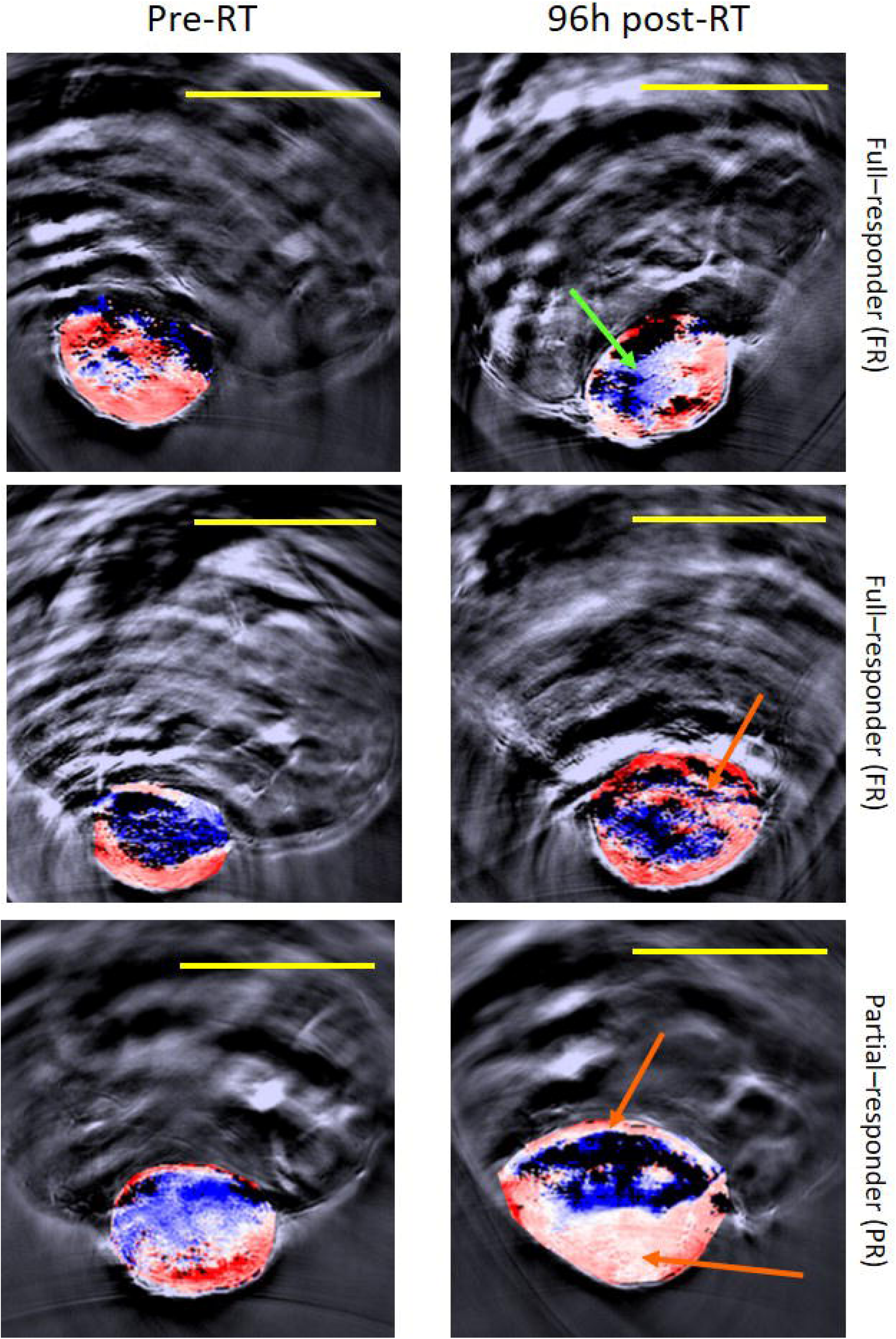
Three representative examples of changes in blood sO_2_ observed 96 h post-RT compared to pre-RT for the 30 Gy cohort (air breathing). Yellow bars = 10 mm. Green arrow indicates regions with decreased blood sO_2_ 96 h post-RT, white to blue regions, comparatively to the pre-RT imaging in one FR. Orange arrows indicate regions with increased blood sO_2_ 96 h post-RT comparatively to pre-RT imaging, in one FR and PR tumours.

The percentage change in blood sO_2_ at 24 and 96 h post 30 Gy irradiation with respect to the baseline blood sO_2_, pre-RT, is shown in Fig. 8 and Table 2. In contrast to the FRs of the 10 Gy and 20 Gy cohort, the FRs of the 30 Gy cohort, demonstrated a mean positive percentage change in blood sO_2_ at both time-points 24 h (6±17% and 6±11% for air-and oxygen-breathing, respectively) and 96 h (7±18% and 13±18% for air-and oxygen-breathing, respectively), post RT. Of the full-responders, 5/8 animals showed an increase in sO_2_ during air-breathing (19±10%) and 6/8 animals during oxygen-breathing (22±11%), 96 h post-RT. Similar to the 10 Gy and 20 Gy cohort, the PRs of the 30 Gy cohort demonstrated a mean positive percentage change (>15±2%) in blood sO_2_ for both time points. During oxygen-breathing, tumours showing a full response had significantly higher blood sO_2_ than controls at 96 h post-RT, whilst the blood sO_2_ of partial-responders was significantly higher at both 24 h and 96 h post-RT.

**Figure 8:**
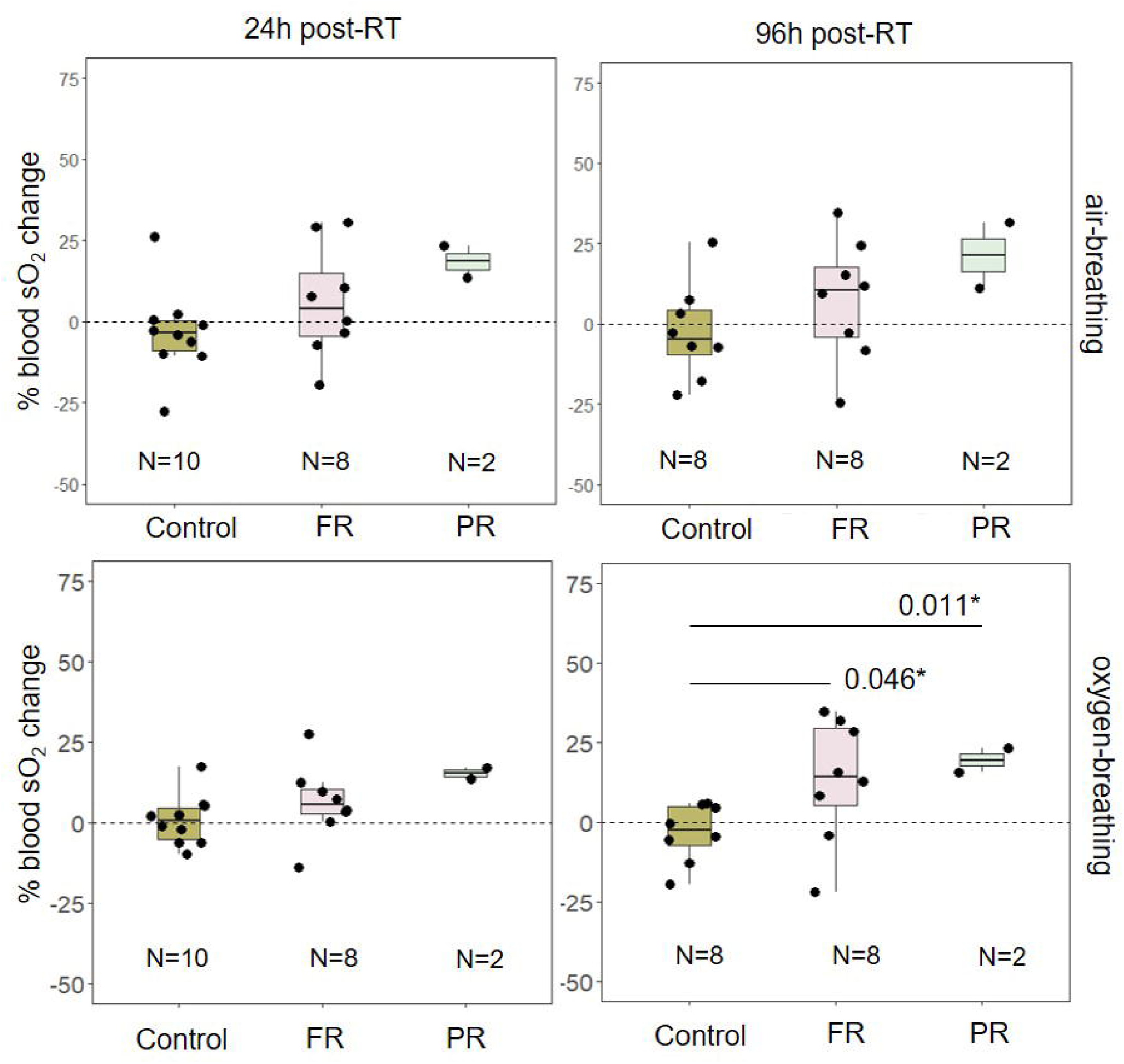
Percentage change in average blood sO_2_ for the 30 Gy RT group between 24 hours, left column, and 96 hours post-RT, right column, and immediately pre-irradiation. Black circles represent the change in average blood sO_2_ for each tumour measured over 3 central slices. Black horizontal lines within the boxplots represent median average blood sO_2_ change per group, with the box extremities representing the 25% and 75% percentiles and the whiskers representing the maximum and minimum values per group. FR = full-responders; PR = partial-responders. * p-value < 0.05; ** p-value < 0.01 using a paired Student’s t-test.

#### *ΔsO*_*2*_ *and haemoglobin results*

The differences in ΔsO_2_ between immediately pre-RT and the two time points after treatments, 24 h and 96 h, were also investigated. No significant differences or trends for ΔsO_2_ were found between treatment response groups, for any of the dose cohorts (10, 20 and 30 Gy), at either 24 h or 96 h post-RT (Tables S4-S6).

Tumour haemoglobin levels 24 h and 96 h post-RT were not statistically significantly different from baseline for any of the treatment response (FR, PR and NR) or control groups, regardless of RT dose (10, 20 and 30 Gy; Tables S4-S6). One exception, however, was that 2 PR tumours, irradiated with 30 Gy, had a significant decrease of 65% or 44% in the level of deoxy-haemoglobin (Hb) at 96 h post-RT compared to immediately pre-RT, during air-breathing.

## Discussion

In this study, we have tested the hypothesis that ‘PAI can predict and monitor tumour response to RT’. The tumoural blood sO_2_, ΔsO_2_ and haemoglobin levels (specifically Hb, HbO_2_ and HbT) were estimated using PAI, prior to and shortly after delivering RT doses of 10 Gy, 20 Gy and 30 Gy to test the hypothesis.

Radiotherapy response categories were chosen to reflect the different types of tumour growth behaviour seen in response to radiation treatments. From Fig.1, it is possible to observe that while for 10 Gy, only 3/13 tumours regressed, the majority of tumours treated with 20 Gy (7/10) and 30 Gy (8/10) did so. The regression took longer in the 10 Gy cohort (29±8 days) than in the 20 Gy (9±6 days) and 30 Gy (11±5 days) groups. This was expected as the higher the ionising dose, the greater the cellular damage, with more tumour cells dying [30, 31]. The proportion of NR was greater for 10 Gy (5/13) than for 20 Gy (2/10), and all 30 Gy treatments resulted in full-or partial-response.

The PR group contained both tumours that regrew and those that stopped growing after RT, with no tumour regression. One of the consequences of hypoxia is that cells enter a dormant state, in which they proliferate more slowly than normoxic cells, but also remain viable for long periods of time [32]. If new vasculature is formed or reoxygenation occurs after radiotherapy, these cells may leave their dormant state and proliferate [33]. This may be the mechanism involved in the different behaviour of PR tumours seen after RT, with tumours that regrew possibly being reoxygenated during the 60 day follow up period of this study, while tumours with growth inhibition were not.

Hypoxia is a microenvironmental feature which results in radioresistance and, consequently, increases the risk of unsuccessful RT treatment. This study has investigated whether tumour blood sO_2_ measured with PAI, 24 h or immediately before treatment, could predict response to 10, 20 or 30 Gy irradiation doses. The results showed a clear trend for FRs having higher average pre-treatment blood sO_2_, in the 10 and 20 Gy radiation groups, than the PRs or NRs (Figs. 2 and 3). The FR tumours from the 10 Gy cohort also had significantly higher baseline blood sO_2_ values (Fig. 2) than the controls, suggesting that high pre-treatment sO_2_ (>0.65) might be predictive of a good response to radiotherapy.

In line with this, blood sO_2_ levels at baseline of 10 and 20 Gy non-/partial-responders were significantly lower than those of the controls (Figs. 2 and 3). This indicates that tumours with relatively low blood sO_2_ might respond poorly to treatment. Nevertheless, while for 10 Gy only a partial-response was observed, for the 20 Gy cohort a full-response was obtained, probably due to the greater level of radiation damage to the tumour. The results for the 30 Gy group show a full response to treatment for a wider initial sO_2_ range (0.38 to 0.68 pre-RT for air-breathing), and hence an increased percentage of tumour remissions, 80%, was observed (Figs. 1 and 4). Since 30 Gy is a high single fraction dose, it is likely that even hypoxic cells are not resistant to the dose. This treatment efficacy was also seen for one animal in the 20 Gy radiation group which had a low sO_2_ value of 0.43±0.02 immediately pre-RT.

The potential to use PAI in the clinic to screen patients before radiotherapy and thus to personalise their radiotherapy exposure to their measured blood sO_2_ would allow the reduction of dose in some patients to avoid side effects and its increase in others to avoid reduced efficacy. Other research groups have demonstrated, in clinical and pre-clinical models, that tumour oxygenation levels prior to radiotherapy correlate with treatment outcome [34, 35]. For example, Nordsmark et al. [36], in 1996, found that patients’ head and neck tumours with low (< 2.5 mmHg) partial oxygen (pO_2_), measured using oxygen electrodes, had significantly lower loco-regional tumour control. Similarly, a better RT outcome for better oxygenated tumours (pO_2_ > 2.5 mmHg) has been shown in subcutaneous C3H mammary carcinomas, in mice [37]. HÖckel et al. [38], using needle electrodes, found that cervical cancer patients with low tumour pO_2_ (median threshold <10 mmHg) tumours had significantly worse overall survival than those with better oxygenated tumours (80 months). Non-invasive imaging techniques, such as positron emission tomography (PET) [37, 39-41] and magnetic resonance imaging (MRI) [42-45] have also been used before delivering RT to infer hypoxia, demonstrating that tumours with higher hypoxic fractions were associated with worse response to treatment than those with lower hypoxic fraction, similar to our findings. To the authors’ knowledge, there are no studies which compare the capability of PAI, MRI and PET for predicting RT effectiveness but this would be worth investigating. Besides the analysis of blood sO_2_, the individual components of haemoblogin, deoxy-(Hb) and oxyhaemoglobin (HbO_2_), as well as the total amount of haemoglobin (HbT), were analysed to investigate their relationship with treatment outcome. The total haemoglobin findings were concordant with those obtained in another study by Rich et al. [25], suggesting this parameter is not a predictive factor for tumour response to radiotherapy. Nordsmark and Overgaard [46], while investigating the relationship between hypoxia and tumour control after radiotherapy, showed that oxygenation in tumours, measured using Eppendorf oxygen electrodes, was independent of total haemoglobin levels, obtained from a venous blood sample acquired before the treatment. Also, Joseph et al. [47] found that a higher coefficient of variation (CoV) was expected for haemoglobin than for sO_2_ over either short periods of time (hours) or longer ones (days), as we have confirmed in a previous study [48]. This higher CoV makes Hb, HbO_2_ and HbT less predictive parameters than blood sO_2_.

Another parameter which was not predictive of treatment outcome was ΔsO_2_. Tomaszwwski et al. [29] has shown in 8 nude mice bearing PC3 prostate cancer that ΔsO_2_ is linearly correlates with an increased uptake of indocyanine green (ICG), suggesting also an increase in HbT. This implies that if haemoglobin was not a predictive factor in radiotherapy, ΔsO_2_ should also not be one.

The differences between average blood sO_2_ before and after treatment were also analysed. In the control cohort, whilst 3 tumours showed an increase in oxygenation (of 3 to 25%, air-breathing) from baseline to 96h post-RT, for all the others (n=7) the opposite effect (-3 to -22%, air-breathing) was observed. It is likely that in the latter cases, tumours were outgrowing their blood supply, whereas in the former we hypothesise that neoangiogenesis (the production of new vasculature) has occurred and oxygenation has met or exceeded the demands of these tumours. In fact, we have shown [28] that tumours with faster growth rates have a larger decrease in blood sO_2_ over time compared to those with lower growth rates.

It is important to highlight that radiation dose, along with type of tumour and level of oxygenation pre-treatment, can affect the level of blood oxygenation post radiotherapy [49]. It has been shown that while conventionally fractionated low-dose radiotherapy (∼2 Gy/fraction) tends to preserve vasculature, hypofractionated-doses (>10 Gy/fraction) are likely to cause vascular collapse, due to endothelial cell death [50, 51], and subsequently a decreased blood sO_2_. This is one of the arguments for using hypofractionated RT, as vascular collapse results in a decrease in oxygen and nutrients to the tumour and consequently in secondary cell death [49]. Garcia-Barros et al. [52], for example, showed that for large single doses (10 to 20 Gy), microvascular damage was observed in murine fibrosarcoma (MCA/129) and melanoma (B16F1) murine xenograft models within 6 hours of irradiation with subsequent tumour cell death.

In our study, for both 10 and 20 Gy exposure groups, i.e. hypofractionated regimes, the FR cohort showed no change, or a mean decrease, in blood sO_2_ both 24 h (0.3 ± 5 to -10 ± 6%) and 96 h post-RT (-8 ± 18 to -13 ± 10%). One possible explanation for this trend is endothelial cell death causing blood vessel collapse. Interestingly, Rich et al. [53] used patient-derived xenograft head and neck squamous cell carcinoma (HNSCC) models and showed a positive correlation between increased PAI measured blood sO_2_ 24 h post-RT (15 Gy) and tumour growth inhibition 2 weeks later. We had similar observations for the FR tumours of the 30 Gy cohort. In contrast to the 10 and 20 Gy cohorts, the FRs in the 30 Gy group had a mean increase in blood sO_2_ at 24 h (6±17% and 6±11%, for air-and oxygen-breathing, respectively) and 96 h post RT (7±18% and 13±18%, for air-and oxygen-breathing, respectively). It can be speculated that for a high dose (30 Gy), the CAL^R^ tumour cells are drastically damaged reducing the oxygen demand. The increase in blood sO_2_ might also be due to acute inflammation arising due to a large radiation dose being delivered in one single fraction [49].

On the other hand, a trend for an increase in post-RT average blood sO_2_ relative to baseline (13 ± 8 to 26 ± 15 %) was observed for PR and NR. This could be due to a lack of endothelial cell death, which then cannot counteract physiological phenomena such as inflammatory processes due to acute radiation treatment or the development of a new matured vascular network after radiation. Both processes are common after radiation and can influence tumour oxygenation [54, 55]. In fact, some studies have suggested that formation of new vessels post-RT is often observed before a tumour regrows and drastically affects tumour control [56-59].

The lack of histological evidence here allows only speculation about the underlying physiological mechanisms occurring after radiotherapy, such as endothelial cell death, tumour reoxygenation or inflammation. Histopathology was not possible for this study as the main aim was to obtain long-term (60 days) tumour volume measurements for the accurate evaluation of response to treatment. In the future, animals will be imaged at regular intervals within the first 24-96 hrs, and oxygenation changes will be evaluated histologically to investigate the timing of these changes. It was also not possible to validate the hypoxic state of tumours before treatment, which could be done using exogenous markers, such as pimonidazole, or other imaging techniques, such as BOLD-MRI or PET. However, our group has shown the good agreement between PAI measurements of blood sO_2_ and pimonidazole staining in another paper [28]. While initial results are encouraging, further investigation is necessary with a larger number of animals, as some radiation exposures resulted in response groups with a small number of animals (<5).

This study shows the potential for using the average blood sO_2_ measured in tumours by PAI, under both medical air (21% O_2_) and 100% oxygen-breathing conditions, before RT to predict their response to radiotherapy, which has been demonstrated here for the first time. In addition, the potential to monitor treatment outcome was demonstrated. PAI measured changes in average blood sO_2_ at 24 h and 96 h post-RT compared to those pre-RT showed a trend for a decrease in this parameter to be correlated with better tumour response, particularly for the later time point. Thus the potential for using photoacoustic imaging as both a predictive and a treatment assessment tool for radiotherapy has been showed in this paper.

## Methods

### Cell line and xenograft model

A human HNSCC cell line, CAL^R^, was cultured in vitro as described previously [60]. Xenografts were established by injecting 5 × 10^5^ cells subcutaneously, into the right flank of female mice (FOXnu^n1^, 6 weeks old, ∼25 grams). All research was conducted under the Guidelines for Animal Welfare as per recommendation by the UK Home Office Animals (Scientific Procedures) Act 1986 [61] and the ARRIVE guidelines [62].

Tumour size was measured non-invasively using callipers every 2 or 3 days, in 3 orthogonal directions (length (l), width (w) and height (h)). An ellipsoidal shape was assumed and the tumour volume (V) calculated using equation (1).

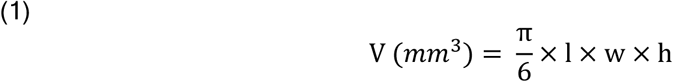

Tumours were measured from 8 to 10 days after cell injection until either the tumour approached the maximum permissible dimensions (average of the 3 dimensions ≤ 12 mm or one dimension reached 15 mm) or 60 days post-radiation was reached.

### Radiation treatment

Animals were irradiated in a computed tomography (CT) imaging guided Small Animal Radiation Research Platform (SARRP, XStrahl, Camberley, United Kingdom). The dose-rate of the SARRP was 2.5 Gy.min^-1^ and energy was 225 kV. Treatment planning was performed using the Muriplan© software provided by the manufacturer [63]. The radiation dose was prescribed to the treatment isocentre which was positioned at the centre of the tumour as visualised in pre-treatment CT images. A square 10×10 mm collimator, the largest available, was chosen to ensure the exposure of the whole tumour. The number and orientation of treatment beams were then chosen. For 30/33 animals, two parallel-opposed beams were used, at 0^°^ and 180^°^ (i.e. vertically downwards and vertically upwards), to obtain a uniform dose distribution within the tumours and to spare adjacent normal tissue. The planning software displays the 10, 20, 60, 80, 95 and 105% isodose lines within the collimator boundaries. For treatment planning the whole tumour was required to lie within the 95% isodose contour. If the tumour was wider than the collimated field, third and fourth beams were added, delivered both at 0^°^ and 180^°^, using a 3 mm or 5 mm collimator to ensure treatment of the whole tumour. For 3 animals, in order to avoid exposure of healthy tissue such as the spinal cord, the treatment delivery angles were adjusted to be oblique i.e. up to 25^°^ and 155^°^. No other techniques for avoiding the exposure of healthy tissue were employed.

When tumour volume reached approximately 200 mm^3^, animals were divided into 4 groups, with each prescribed a different radiation dose which was delivered in a single fraction:

- Controls (0 Gy): N = 10;
- 10 Gy: n=13, total treatment time ∼4 minutes;
- 20 Gy: n=10, total treatment time ∼8 minutes;
- 30 Gy: n=10, total treatment time ∼12 minutes.

### Treatment response assessment

Radiation treatment outcome was evaluated for up to 60 days after delivery, using calipers for tumour volume measurement, and it was classified as follows:

- No response (NR): the rate of tumour growth remains within the range of the control, non-irradiated group with no increase in survival.
- Partial response (PR): either a reduced tumour growth rate following RT or tumour regression after RT followed by regrowth at a later time.
- Full response (FR): tumour size decreased with some tumours becoming undetectable.

### Photoacoustic Imaging

Animals were imaged using the MultiSpectral Optical Tomography small animal imaging device (MSOT, iThera, Germany), as described in [48]. Before imaging, animals were anaesthetised by intraperitoneal injection of a combination of hypnovel (1:3), fentanyl (1:3) and medetomidine (1:3). For all imaging sessions, animals were equilibrated for 5 minutes in water at 34^°^C prior to imaging. Initially, medical air (21%-oxygen) was provided. Immediately after acquisition of the first image dataset in which images of the entire tumour spaced 1 mm apart were obtained, the gas supply was changed to 100% oxygen, which can work as a contrast agent, and a second imaging sequence was started 2 minutes later in order to provide enough time for the oxygen to be distributed throughout the animal. For each image slice, the following optical excitation wavelengths were used: 700, 715, 730, 750, 760, 800, 850 and 900 nm. For each wavelength, photoacoustic signals from 10 pulses were averaged. A model-based inversion algorithm [64] (ViewMsot v3.8, iThera Medical) was used to reconstruct the photoacoustic signal. Neglecting any position or wavelength dependent laser fluence variation, the reconstructed data was spectrally unmixed using a linear regression algorithm provided by the company, to estimate the proportion of oxy-and deoxy-haemoglobin for each image pixel, [25, 64, 65].

For imaging analysis, three central slices 1 mm apart were chosen, based on their distance from tumours edges observed in the reconstructed photoacoustic images. Using the viewMSOT software, the average Hb and HbO_2_ values for each ROI of the subcutaneous tumour were obtained and used to calculate the average HbT, resulting from the addition of both parameters. The average sO_2_ of each tumour slice was calculated as the ratio of average HbO_2_ and the average HbT. For each tumour, the Hb, HbO_2_, HbT and sO_2_ values of the 3 ROIs were averaged. The sO_2_ based on the Hb and HbO_2_ measurements was also calculated on a pixel-by-pixel basis, producing a blood sO_2_ distribution map, or ‘oxy-map’, for each slice. The difference in average blood sO_2_ between air-and oxygen-breathing, ΔsO_2_, was also calculated.

Mice were imaged both 24 hours (-24 h) and immediately before (pre-RT) treatment. Two pre-radiation time points were chosen rather than just one due to the intrinsic variation in tumour oxygenation described in the Introduction. The time points were kept within 24 hours to avoid a significant tumour volume changes. Images were also acquired at two post-treatment time points, 24 hours (+24 h) and 96 hours (+96 h).

### Statistical analysis

A paired 2-tailed Student’s t-test was used to test for significance in all data analysis (R software, version 3.4), after confirming data normality using the Shapiro-Wilk test. Statistical significant (*) was assumed for p-values below 0.05, very significant (**) when p-value was below 0.01 and extremely significant (***) when p-value was below 0.001.

## Acknowledgements

We gratefully thank the Portuguese Foundation for Science and Technology (PhD grant SFRH/BD/90250/2012) for funding part of this project. We further acknowledge funding from the EPSRC (strategic equipment grant EP/NO15266/1) and the Cancer Research UK Cancer Imaging Centre at the Institute of Cancer Research (grant C1060/A16464). We would also like to thank the Focused Ultrasound Foundation for their support of our Centre of Excellence.

## Additional information

### Author contributions

M.M.C., A.S., I.R., G.t.H. and J.B. conceived the experiments, M.M.C. and A.S. conducted the experiments, M.M.C., A.S., I.R., G.t.H. and J.B. analysed the data. T.O helped planning the radiotherapy experiments. C.B. helped with the selection of the appropriate tumour model and provided the CAL^R^ cells. All authors reviewed the manuscript.

### Availability of materials and data

All data is available for the readers upon request.

## References

1. Yoshimura, M., et al., Microenvironment and radiation therapy. Biomed Res Int, 2013. 2013: p. 685308.

2. Barker, H.E., et al., The tumour microenvironment after radiotherapy: mechanisms of resistance and recurrence. Nat Rev Cancer, 2015. 15(7): p. 409–25.

3. Rockwell, S., et al., Hypoxia and radiation therapy: past history, ongoing research, and future promise. Curr Mol Med, 2009. 9(4): p. 442–58.

4. Semenza, G.L., Intratumoral hypoxia, radiation resistance, and HIF-1. Cancer Cell, 2004. 5(5): p. 405–6.

5. Fenner, A., Prostate cancer: Hypoxia predicts relapse and recurrence after radiotherapy. Nat Rev Urol, 2012. 9(5): p. 237.

6. Harada, H., [Hypoxia-inducible factors as molecular targets for cancer therapy]. Nihon Rinsho, 2012. 70 Suppl 8: p. 113–7.

7. Hicks, R.J., Learning from Failure; Hypoxia Is an Evil Foe. J Nucl Med, 2017. 58(7): p. 1043–1044.

8. Brown, J.M. and W.R. Wilson, Exploiting tumour hypoxia in cancer treatment. Nat Rev Cancer, 2004. 4(6): p. 437–47.

9. Joiner, M. and A. Kogel, Basic Clinical Radiobiology (book) 4th edition. Macmillian Publishing Solutions, 2008.

10. Sun, X., et al., Tumor hypoxia imaging. Mol Imaging Biol, 2011. 13(3): p. 399–410.

11. Lopci, E., et al., PET radiopharmaceuticals for imaging of tumor hypoxia: a review of the evidence. Am J Nucl Med Mol Imaging, 2014. 4(4): p. 365–84.

12. Lopci, E., et al., Imaging acute spinal myelitis with 18F-FDG PET/CT. Eur J Nucl Med Mol Imaging, 2014. 41(2): p. 399–400.

13. Ziemer, L.S., et al., Oxygen distribution in murine tumors: characterization using oxygen-dependent quenching of phosphorescence. J Appl Physiol (1985), 2005. 98(4): p. 1503–10.

14. Elas, M., et al., Electron paramagnetic resonance oxygen image hypoxic fraction plus radiation dose strongly correlates with tumor cure in FSa fibrosarcomas. Int J Radiat Oncol Biol Phys, 2008. 71(2): p. 542–9.

15. Imai, T., et al., Direct measurement of hypoxia in a xenograft multiple myeloma model by optical-resolution photoacoustic microscopy. Cancer Biol Ther, 2017. 18(2): p. 101–105.

16. Buxton, R.B., The physics of functional magnetic resonance imaging (fMRI). Rep Prog Phys, 2013. 76(9): p. 096601.

17. Ueda, S., et al., Baseline tumor oxygen saturation correlates with a pathologic complete response in breast cancer patients undergoing neoadjuvant chemotherapy. Cancer Res, 2012. 72(17): p. 4318–28.

18. Bar-Zion, A., et al., Functional Flow Patterns and Static Blood Pooling in Tumors Revealed by Combined Contrast-Enhanced Ultrasound and Photoacoustic Imaging. Cancer Res, 2016. 76(15): p. 4320–31.

19. Gerling, M., et al., Real-time assessment of tissue hypoxia in vivo with combined photoacoustics and high-frequency ultrasound. Theranostics, 2014. 4(6): p. 604–13.

20. Hardee, M.E., et al., Novel imaging provides new insights into mechanisms of oxygen transport in tumors. Curr Mol Med, 2009. 9(4): p. 435–41.

21. Beard, P., Biomedical photoacoustic imaging. Interface Focus, 2011. 1(4): p. 602–31.

22. Zhang, H.F., K. Maslov, and L.V. Wang, In vivo imaging of subcutaneous structures using functional photoacoustic microscopy. Nat Protoc, 2007. 2(4): p. 797–804.

23. Raes, F., et al., High Resolution Ultrasound and Photoacoustic Imaging of Orthotopic Lung Cancer in Mice: New Perspectives for Onco-Pharmacology. PLoS One, 2016. 11(4): p. e0153532.

24. Rich, L.J. and M. Seshadri, Photoacoustic imaging of vascular hemodynamics: validation with blood oxygenation level-dependent MR imaging. Radiology, 2015. 275(1): p. 110–8.

25. Tzoumas, S., et al., Eigenspectra optoacoustic tomography achieves quantitative blood oxygenation imaging deep in tissues. Nat Commun, 2016. 7: p. 12121.

26. Burton, N.C., et al., Multispectral opto-acoustic tomography (MSOT) of the brain and glioblastoma characterization. Neuroimage, 2013. 65: p. 522–8.

27. Kruger, R.A., et al., Photoacoustic angiography of the breast. Med Phys, 2010. 37(11): p. 6096–100.

28. Martinho Costa, M., et al., Quantitative photoacoustic imaging study of tumours in vivo: baseline variations in quantitative measurements. bioRxiv, 2018.

29. Tomaszewski, M.R., et al., Oxygen Enhanced Optoacoustic Tomography (OE-OT) Reveals Vascular Dynamics in Murine Models of Prostate Cancer. Theranostics, 2017. 7(11): p. 2900–2913.

30. Brown, J.M., The hypoxic cell: a target for selective cancer therapy-- eighteenth Bruce F. Cain Memorial Award lecture. Cancer Res, 1999. 59(23): p. 5863–70.

31. Shinomiya, N., New concepts in radiation-induced apoptosis: ’premitotic apoptosis’ and ’postmitotic apoptosis’. J Cell Mol Med, 2001. 5(3): p. 240–53.

32. Hayat, M.A., Tumor dormancy, quiescence, and senescence. aging, cancer, and noncancer pathologies Volume 3 Volume 3. 2014.

33. Hoppe-Seyler, K., et al., Induction of dormancy in hypoxic human papillomavirus-positive cancer cells. Proc Natl Acad Sci U S A, 2017. 114(6): p. E990–E998.

34. Colliez, F., B. Gallez, and B.F. Jordan, Assessing Tumor Oxygenation for Predicting Outcome in Radiation Oncology: A Review of Studies Correlating Tumor Hypoxic Status and Outcome in the Preclinical and Clinical Settings. Frontiers in Oncology, 2017. 7.

35. Park, H.J., et al., Radiation-Induced Vascular Damage in Tumors: Implications of Vascular Damage in Ablative Hypofractionated Radiotherapy (SBRT and SRS). Radiation Research, 2012. 177(3): p. 311–327.

36. Nordsmark, M., M. Overgaard, and J. Overgaard, Pretreatment oxygenation predicts radiation response in advanced squamous cell carcinoma of the head and neck. Radiotherapy and Oncology, 1996. 41(1): p. 31–39.

37. Mortensen, L.S., et al., Accessing radiation response using hypoxia PET imaging and oxygen sensitive electrodes: A preclinical study. Radiotherapy and Oncology, 2011. 99(3): p. 418–423.

38. Hockel, M., et al., Association between tumor hypoxia and malignant progression in advanced cancer of the uterine cervix. Cancer Research, 1996. 56(19): p. 4509–4515.

39. Schutze, C., et al., Effect of [F-18]FMISO stratified dose-escalation on local control in FaDu hSCC in nude mice. Radiotherapy and Oncology, 2014. 111(1): p. 81–87.

40. Tran, L.B.A., et al., Potential role of hypoxia imaging using F-18-FAZA PET to guide hypoxia-driven interventions (carbogen breathing or dose escalation) in radiation therapy. Radiotherapy and Oncology, 2014. 113(2): p. 204–209.

41. Mortensen, L.S., et al., FAZA PET/CT hypoxia imaging in patients with squamous cell carcinoma of the head and neck treated with radiotherapy: Results from the DAHANCA 24 trial. Radiotherapy and Oncology, 2012. 105(1): p. 14–20.

42. Ovrebo, K.M., et al., Assessment of Tumor Radioresponsiveness and Metastatic Potential by Dynamic Contrast-Enhanced Magnetic Resonance Imaging. International Journal of Radiation Oncology Biology Physics, 2011. 81(1): p. 255–261.

43. Ellingsen, C., et al., DCE-MRI of the hypoxic fraction, radioresponsiveness, and metastatic propensity of cervical carcinoma xenografts. Radiotherapy and Oncology, 2014. 110(2): p. 335–341.

44. Rodrigues, L.M., et al., Tumor R-2 * is a prognostic indicator of acute radiotherapeutic response in rodent tumors. Journal of Magnetic Resonance Imaging, 2004. 19(4): p. 482–488.

45. Shukla-Dave, A., et al., Dynamic Contrast-Enhanced Magnetic Resonance Imaging as a Predictor of Outcome in Head-and-Neck Squamous Cell Carcinoma Patients with Nodal Metastases. International Journal of Radiation Oncology Biology Physics, 2012. 82(5): p. 1837–1844.

46. Nordsmark, M. and J. Overgaard, Tumor hypoxia is independent of hemoglobin and prognostic for loco-regional tumor control after primary radiotherapy in advanced head and neck cancer. Acta Oncol, 2004. 43(4): p. 396–403.

47. Joseph, J., et al., Evaluation of Precision in Optoacoustic Tomography for Preclinical Imaging in Living Subjects. J Nucl Med, 2017. 58(5): p. 807–814.

48. Martinho Costa, M., et al., Quantitative photoacoustic imaging study of tumours in vivo. bioRxiv, 2018.

49. Park, H.J., et al., Radiation-induced vascular damage in tumors: implications of vascular damage in ablative hypofractionated radiotherapy (SBRT and SRS). Radiat Res, 2012. 177(3): p. 311–27.

50. Jani, A., et al., High-Dose, Single-Fraction Irradiation Rapidly Reduces Tumor Vasculature and Perfusion in a Xenograft Model of Neuroblastoma. Int J Radiat Oncol Biol Phys, 2016. 94(5): p. 1173–80.

51. Song, C.W., et al., Radiobiological basis of SBRT and SRS. Int J Clin Oncol, 2014. 19(4): p. 570–8.

52. Garcia-Barros, M., et al., Tumor response to radiotherapy regulated by endothelial cell apoptosis. Science, 2003. 300(5622): p. 1155–9.

53. Rich, L.J. and M. Seshadri, Photoacoustic monitoring of tumor and normal tissue response to radiation. Sci Rep, 2016. 6: p. 21237.

54. Kozin, S.V., et al., Neovascularization after irradiation: what is the source of newly formed vessels in recurring tumors? J Natl Cancer Inst, 2012. 104(12): p. 899–905.

55. Shaked, Y. and E.E. Voest, Bone marrow derived cells in tumor angiogenesis and growth: are they the good, the bad or the evil? Biochim Biophys Acta, 2009. 1796(1): p. 1–4.

56. Solesvik, O.V., E.K. Rofstad, and T. Brustad, Vascular Changes in a Human-Malignant Melanoma Xenograft Following Single-Dose Irradiation. Radiation Research, 1984. 98(1): p. 115–128.

57. Kioi, M., et al., Inhibition of vasculogenesis, but not angiogenesis, prevents the recurrence of glioblastoma after irradiation in mice. Journal of Clinical Investigation, 2010. 120(3): p. 694–705.

58. Song, C.W. and S.H. Levitt, Vascular Changes in Walker-256 Carcinoma of Rats Following X-Irradiation. Radiology, 1971. 100(2): p. 397-&.

59. Denis, F., et al., Radiosensitivity of rat mammary tumors correlates with early vessel changes assessed by power Doppler sonography. Journal of Ultrasound in Medicine, 2003. 22(9): p. 921–929.

60. Baker, L.C.J., et al., Evaluating imaging biomarkers of acquired resistance to targeted EGFR therapy in xenograft models of human squamous cell carcinoma of the head and neck (SCCHN). Cancer Research, 2013. 73(8).

61. Hollands, C., The Animals (Scientific Procedures) Act 1986. Lancet, 1986. 2(8497): p. 32–33.

62. Drummond, G.B., D.J. Paterson, and J.C. McGrath, ARRIVE: new guidelines for reporting animal research. Experimental Physiology, 2010. 95(8): p. 841–841.

63. Jacques, R., et al., Real-time dose computation: GPU-accelerated source modeling and superposition/convolution. Med Phys, 2011. 38(1): p. 294–305.

64. Rosenthal, A., et al., Fast semi-analytical acoustic inversion for quantitative optoacoustic tomography. Photons Plus Ultrasound: Imaging and Sensing 2011, 2011. 7899.

65. Rosenthal, A., V. Ntziachristos, and D. Razansky, Acoustic Inversion in Optoacoustic Tomography: A Review. Current Medical Imaging Reviews, 2013. 9(4): p. 318–336.

